# Extensive horizontal exchange of transposable elements in the *Drosophila pseudoobscura* group

**DOI:** 10.1101/284117

**Authors:** Tom Hill, Andrea J. Betancourt

## Abstract

While the horizontal transfer of a parasitic element can be a potentially catastrophic, it is increasingly recognized as a common occurrence. The horizontal exchange, or lack of exchange, of TE content between species results in different levels of divergence among a species group in the mobile component of their genomes. Here, we examine differences in the TE content of the *Drosophila pseudoobscura* species group. We identify several putative horizontal transfer events, and examine the role that horizontal transfer plays in the spread of TE families to new species and the homogenization of TE content in these species. Despite rampant exchange of TE families between species, we find that both TE content differs hugely across the group, likely due to differing activity of each TE family and differing suppression of TEs due to divergence in Y chromosome size, and its resulting effects of TE regulation. Overall, we show that TE content is highly dynamic in this species group, and that it plays a large role in shaping the differences seen between species.

**Data availability:** All data used in this study (summarized in table S1) is freely available online through the NCBI short read archive (NCBI SRA: ERR127385, SRR330416, SRR330418, SRR1925723, SRR330426, SRR330420, SRR330423, SRR617430-74). All genomes used are either available through *flybase*.*org* or *popoolation*.*at*.

## Introduction

Unlike mammals, which have few active transposable elements (TEs) mostly fixed insertions within species (Hellen and Brookfield 2013a; b), transposable elements (TEs) in *Drosophila* species appear to be highly active, as inferred from a high proportion of polymorphic, and thus presumably recent, insertions (Charlesworth and Langley 1989; Sniegowski and Charlesworth 1994; Charlesworth *et al*. 1997; González *et al*. 2008; Petrov *et al*. 2011). The dynamic nature of TEs in *Drosophila* is reflected in the data from the 12-genomes project (Clark *et al*. 2007). While species in the genus all host LTR, non-LTR retrotransposons and TIR DNA transposons in roughly the same rank order of abundance (Sessegolo *et al*. 2016), the contribution of each appears to differ for different genomes. The proportion of total TE content that is non-LTRs, for example, ranges from ∼12% to ∼35% (Clark *et al*. 2007; Sessegolo *et al*. 2016).

Under a model of TE evolution where active transposition is followed by suppression and eventually, inactive decayed elements, one might expect that the active families of elements would differ between the species (Kaplan *et al*. 1985; Maruyama and Hartl 1991; Capy *et al*. 1992; Lohe *et al*. 1995; Hartl *et al*. 1997). Instead, the overall content is largely similar (Vieira *et al*. 1999; Lerat *et al*. 2011; Kofler *et al*. 2015b), with many of the same TE insertions found at low frequencies in both species (Kofler *et al*. 2015b). The reason might be that the overall TE content between species be regularly homogenized by horizontal exchange between the species (Bartolomé *et al*. 2009). This process is exemplified by the recent horizontal transfer of the P-element, newly acquired by *D. melanogaster* sometime in the 20^th^ century from a Caribbean species, into *D. simulans* (Kofler *et al*. 2015a; Hill *et al*. 2016).

Here, we investigate these questions in a different *Drosophila* group, the *pseudoobscura* group, using publicly available genome sequences for five species, and an improved genome sequence from *D. pseudoobscura* (Richards *et al*. 2005), and several sequenced third chromosome isolates (Fuller *et al*. 2016). Unlike *D. simulans* and *D. melanogaster*, these species are not cosmopolitan and thus may have had less opportunity to encounter new transposable elements outside their ancestral range. Further, in contrast to most Drosophila, some species in this group were reported to have mostly fixed insertions; we re-examine this claim with new data. We also examine horizontal exchange between species within the group and from outside the group, and find abundant evidence of recurrent horizontal exchange.

## Materials and Methods

### Sequence data

All sequence data used is summarized in Table S1. We used publicly available reference genomes for five species: *D. pseudoobscura* (NCBI: PRJNA18793), *D. persimilis* (NCBI: PRJNA29989 genome assembled from Sanger sequence reads, http://popoolation.at/persimilis_genome/ for the genome based on illumina reads), *D. affinis* (NCBI: ERX103526), *D. lowei* (http://popoolation.at/lowei_genome/; *Palmieri et al. 2014), D. miranda* (NCBI: PRJNA77213) and *D. affinis* (http://popoolation.at/affinis_genome/). We also used publicly available paired-end illumina data from inbred lines for four of these species [*D. persimilis* (SRA: SRR330426), *D. miranda* (SRA: SRR1925723), *D. lowei* (SRA: SRR330416 and SRR330418) and *D. affinis* (ENA: ERR127385)]. As we were unable to find publicly available paired-end illumina data for *D. pseudoobscura*, we used a data generated from an individual wild *D. pseudoobscura* made homozygous for the reference third chromosome inversion type (SRA: SRR617430, S. Schaeffer, pers. Comm.; Fuller et al. 2016). As a result, only the third chromosome represents a wild chromosome, the rest of the genome is a mosaic of material from the wild and from the two different balancer stocks used, due to this we limited any population statistical analysis to the third chromosome.

### De novo annotation of transposable elements in the D. pseudoobscura group

We annotated TE families in all five species, as well as putative TE sequences in the more diverged species (such as *D. lowei* and *D. affinis*), and compared our *de novo* annotations to the previous annotations for *D. pseudoobscura* and *D. persimilis*. These sequences were identified using *RepeatModeler* and *LTRHarvest* (Ellinghaus *et al*. 2008; Smit and Hubley 2008) and filtered, as outlined in Supplementary Figure 1 to give us a set of ‘high confidence’ TE annotations.

**Figure 1:**
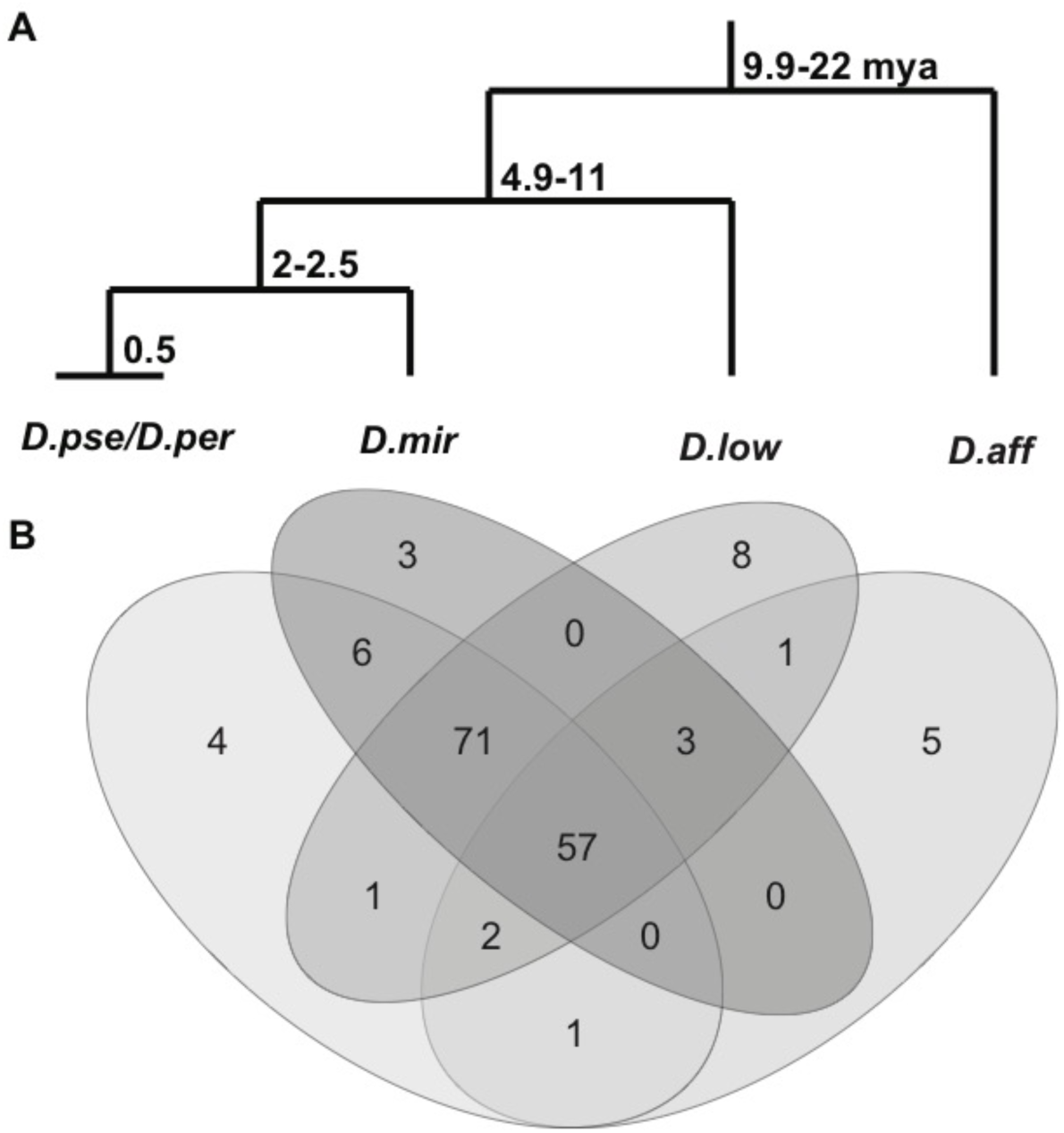
**A**. Phylogeny of the *D. pseudoobscura* group and the estimated time of divergence between nodes. **B**. Number of TE families shared between species in the *D. pseudoobscura* group, including putative novel families.

To *de novo* annotate the transposable elements, as shown in Figure S1:

1. We recovered a set of TE candidates for each species using the reference genomes. We used two separate pipelines: *(i) Repeatscout* and *PILER* in the *RepeatModeler* pipeline (default parameters) (Price *et al*. 2005; Smit and Hubley 2008), with all sequences designated as microsatellites and simple repeats removed from the output, and *(ii) LTRHarvest*, which finds LTR retrotransposons (using parameters recommended in the *LTRHarvest* manual: -tis *-suf* -lcp -d*es -sds –dna;* -seed 10*0* - minlenltr 100 -maxlenltr 1000 -mindistltr 1000 -maxdistltr 15000 -xdrop 5 -mat 2 –mis −2 -ins −3 -del −3 -similar 90.0 -overlaps best -mintsd 5 -maxtsd 20 -motif tgca -motifmis 0 -vic 60 -longoutput) (Ellinghaus *et al*. 2008). Though this step may bias us to find primarily LTRs, we note that most previously known TEs we find are LTRs, while most (19 of 41) novel elements are DNA transposons (Table S2).
2. Step 1 resulted in a set of 769 candidate TE sequences, ranging from 208bp to 14.5kb. We used BLAST to filter and annotate the candidate TEs (parameters: e-value < 1e- 08, -word_size 10, -perc_identity 85) (Altschul *et al*. 1990), by searching a database of all known *Repbase* and *Flybase* transposable element sequences for *Diptera* (including 121 TEs previously found in *D. pseudoobscura, D. persimilis* or *D. miranda*), with sequenced duplicated between the data bases removed using a custom python script.
  a. Sequences that show single BLAST hits (e-value ≤ 1e-08) to this data base were assumed to represent a previously identified TE family. We discarded these sequences and used the Repbase/Flybase TE sequence to represent the family instead. (349 sequences).
  b. From the remaining sequences, those that showed BLAST hits to several TE families, all from one superfamily, were considered to potentially represent a previously unidentified family within that superfamily. (180 sequences).
  c. Of the remaining sequences, those with hits all in a single order, but to multiple superfamilies, were potentially novel TEs within this order. (18 sequences).
  d. For sequences which had no potential TE family assigned in Step 2 (222 sequences), we attempted to find matches by aligning them to the online NCBI non-redundant database using megablast. Of these, 202 had annotated or predicted genes as the primary BLAST hit; these were discarded. The remaining potentially novel TEs were retained (20 sequences),

To facilitate downstream analysis, we obtained a single representative sequence for the potential novel TEs identified in Steps 2b, c and d, as is already done for those in Step 2a. To do this, we clustered sequences found for all species using *vmatch* (recommended *LTRHarvest* parameters: -dbcluster 95 7 -p -d -seedlength 50 -l 1101 -exdrop 9) (Kurtz 2010). We confirmed these clusters by BLASTing novel TE sequences to themselves and grouping them by similar matches (parameters: e-value < 0.00001, -word_size 10).

4. As these may only represent partial TE sequences, we further assembled the grouped sequences using *Trinity* (default parameters) to collapse similar sequences and get a representative sequence for the cluster, even if only a fragment of the consensus sequence (Haas *et al*. 2013). We checked these assemblies and clusters by aligning sequences from the cluster and with the *Trinity* assembly (if applicable) using *MAFFT* (parameters: --thread 3 --threadit 0 –reorder --leavegappyregion –auto) (Katoh *et al*. 2002), to ensure that the assembly or longest sequence representing the putative novel TE was recovered. From each cluster of similar sequences, we took the longest sequence as the representative fragment of each putatively novel family.

5. Some of the putatively novel families identified in 2b may instead be divergent representatives of known families. To see whether this was the case, we again attempted to identify previously known families among them using the consensus sequences from the five species genomes. We aligned novel TEs pairwise to all *Repbase* TEs using *MAFFT* (parameters: --thread 3 --threadit 0 --reorder -- leavegappyregion --auto) and used a custom *python* script to find the number of diverged aligned bases. We defined sequences as belonging to a known family if they were >90% similar to a known family across the sequence, following (Kohany *et al*. 2006). Two families of the novel sequences were found to belong to known families in this way (an I-element and a Jockey element), but were closely related to insertions in distant relatives of the *obscura* group (*I*-4_DF from *D. funebris* and *Jockey*-8_DRh from *D. rhopaloa*, respectively). We therefore retained these sequences in our data set, as they likely represent diverged copies of these families, or ancient horizontal acquisitions.

6. From Steps 1-5, we found 567 candidate TE sequences, 349 of which belong to previously described TE families, including all 121 families previously found in the *pseudoobscura* group (‘known’ families), and 445 others (putative ‘novel’ families). We proceeded to filter sequences from this set which were represented by very few or very short matches to the reference genomes.

a. First, we used the 567 sequences to repeat mask the reference genome of each species using *RepeatMasker* (parameters: –no_is –norna –no_low – gff –gccalc –u –s –cutoff 200) (Tarailo-Graovac and Chen 2009), following recommendations in (Kofler *et al*. 2012). We required that the families have at least 25 Repeatmasker hits in at least one species (237 sequences retained, 116 known and 121 novel families).
b. We then estimated the copy number of each TE family for each species from the Illumina short read data from adult females, discarding those estimated to have a median coverage less than 2-fold that of the third chromosome for less than 80% of the length of the sequence. To do this, we mapped short reads to the repeated masked reference genome and the 237 TE sequences retained from the previous step using BWA MEM (parameters: paired end –t 5 -M) (Li and Durbin 2009), and estimated coverage with *bedtools genomecov* (Quinlan and Hall 2010). Due to the poor assembly of the *D. persimilis* genome, we used a reference consisting of the *D. pseudoobscura* genome and the *D. persimilis* TE sequences. (157 sequences retained, from 116 known and 41 families novel to this species group).

We considered these 157 sequences to be a cromulent representation of the TE content in the *pseudoobscura* group, though we recognize that we may have discarded some true TE sequences.

Using this method, we found strong support for 114 of the 121 TE families previously described in *D. pseudoobscura, D. persimilis* or *D. miranda* and 2 TEs previously identified in other species. We also found 41 novel sequences, including two subfamilies of previously known sequences, 30 newly assembled sequences which BLAST exclusively to one super family, and nine potentially new families that BLAST to one TE order. We also found 15 sequences that cannot be assigned an order (either due to BLAST hits to multiple orders, or no BLAST hits). These 15 sequences passed all filters, including being found multiple times in species genomes and did not correspond to genes or other NCBI sequences in a universal BLAST search. To avoid unreliable inferences, we discarded these sequences from downstream analyses, but gave each of the 41 novel sequences an ID (Table S2), and included them in masking and mapping stages. Sequences are available in File S1.

### Estimating TE density in the reference genome

We used *RepeatMasker* v. 4.0.6 to mask each reference genome using the 157 consensus TE sequences and 15 unknown sequences from the *de novo* annotation, (parameters: –no_is –norna –nolow –gff –gccalc –u –s –cutoff 200) (Tarailo-Graovac and Chen 2009). To estimate TE density, we used the density of TE bases per 1MB sliding window (with a step size of 100kb) of the *D. pseudoobscura* reference genome (after removing all N bases [e.g. TE bases / [window size – Ns in chromosome]]), across both assembled scaffolds and unassembled contigs from the reference genome.

### Identifying insertions in reference genomes and in sequenced third chromosome lines of D. pseudoobscura

To identify insertion sites in the reference genomes of *D. pseudoobscura* and *D. persimilis*, we used the *PopoolationTE2* pipeline (Kofler *et al*. 2016a). Briefly, we used *RepeatMasker* v. 4.0.6 to mask the *pseudoobscura* genome using the 157 consensus TE sequences and 15 unknown sequences identified above (parameters: –no_is –norna – nolow –gff –gccalc –u –s –cutoff 200) (Tarailo-Graovac and Chen 2009). We chose to use the *D. pseudoobscura* reference, rather than the fragmented *D. persimilis* reference, as it facilitated mapping reads to genomic insertion sites. We expect similar results as these species are closely related (0.018 average synonymous divergence (Noor *et al*. 2007)), and we find that similar numbers of reads map to TEs regardless of whether the *D. pseudoobscura* or *D. persimilis* genome is used (27.63% vs 27.27%).

We then mapped available Illumina reads to the repeat masked reference, the consensus TE sequences, and to sequences matching these consensus TEs identified by *RepeatMasker* using BWA-MEM (parameters: paired end –t 5 -M, with secondary alignments reported, but marked) (Li and Durbin 2009). Using masked TE sequences to aids mapping of degenerate TE sequences, as described in (Kofler *et al*. 2016a).

Following mapping, we generated a ppileup file summarizing identities and locations of TE insertions for all lines in *PopoolationTE2* (default settings, --map-qual 10) and subsampled to a physical coverage of 25, removing secondary alignments. As these sequences are mostly from inbred lines, we required the estimated frequency to be at least 50% (default parameters, --target-coverage 25, --min-count 5, minimum frequency = 0.5) (Kofler *et al*. 2016a). We then identified the number of insertions per MB window (after adjusting for the number of N bases in the window [e.g. TE number / [window size – Ns in window]]) across the genome of each species.

### Expression confirmation of putative TE sequences

We also used expression data for mRNA (SRA: SRR1956914, taken from (Duff *et al*. 2015)) and small RNAs (SRA: SRR032435, taken from (Leslie *et al*. 2010)) from the *D. pseudoobscura* reference line (MV-25) to examine the expression of novel TEs.

Before further analysis, we trimmed all genomic and RNAseq Illumina reads used with *Sickle* to remove low quality sequence data (default parameters for long reads, minimum length = 16 for small RNAs), and removed reads that were unpaired (apart from the small RNA reads) after this step from the sequence data (Joshi and Fass 2011).

We mapped small RNA sequences from *D. pseudoobscura* to known and novel TEs identified in that species, using publicly available small RNA reads from the reference strain ((Leslie *et al*. 2010), SRA: SRR032435).

We first removed non-TE related small RNAs, following (Aravin *et al*. 2007; Rahman *et al*. 2015), by mapping to a database of known *Drosophila* viruses and small RNAs other than those that are TE-related, including miRNAs, viral siRNAs, snoRNA (Rahman *et al*. 2015), using *BWA aln* and allowing for up to 3 mismatches (parameters: -n 3) (Aravin *et al*. 2007; Li and Durbin 2009). We then mapped the remaining reads to the repeat masked *D. pseudoobscura* reference genome and the novel and known TE sequences identified in this study (*BWA aln* parameters: -n 3, maximum 2 alignments).

We classified small RNAs by length and orientation using a custom python script and the *Pysam* python library, following (Brennecke *et al*. 2008). Specifically, we considered small RNAs from 21 to 23 to be siRNAs and from 24 to 29 to be piRNAs (Obbard *et al*. 2009). We used *bedtools* (*intersect*, -wa –wb –f 0.3 –r), to check for a 10- bp overlap between sense and anti-sense matches and used *sequence logos* (Schneider and Stephens 1990) to check for the 1-T, 10-A bias, both associated with ping-pong amplification, a characteristic feature of piRNAs (Levine and Malik 2011).

### Detecting short range horizontal transfer events within the pseudoobscura group

To detect horizontal transfer of TEs within the five species examined, we compared divergence between consensus TE sequences to genomic divergence, following the rationale described in (Bartolomé *et al*. 2009). We limited this analysis to families found in at least 3 species and with an annotation on Repbase.

To construct consensus TE sequences for each TE family and each species, we identified the major allele for each species at each variable site using *GATK v3*.*5-0-g36282e4 HaplotypeCaller*, with ploidy levels set to the estimated copy numbers based on coverage of the TE sequence, and using *FastaAlternateReferenceMaker* (default parameters) to generate fasta sequences from the mapped data (DePristo *et al*. 2011).

We aligned these consensus sequences from each species using *MAFFT* (parameters: --thread 3 --threadit 0 --reorder --leavegappyregion –auto) (Katoh *et al*. 2002) and generated a phylogeny of each sequence using the *Repbase* annotation and *PhyML* (parameters: -M GTR) (Guindon *et al*. 2010). We obtained a total of 39 annotated alignments that included sequences for *D. affinis* comparisons, and 62 additional sequences for all other species comparisons (noted in Table S2).

We estimated synonymous site divergence (*d*_*S*_) in the TE sequences pairwise between species using *codeml* (with transition–transversion rates estimated from the data, and codon frequencies from the nucleotide frequencies) and the coding regions for these TEs as annotated in *Repbase* (Kohany *et al*. 2006; Yang 2007). We then compared *d*_*S*_ of TEs to that of orthologous genes between species obtained in the same way, taken from Avila et al. (2014). Following Bartolomé et al. (2009), we considered an individual family to show strong evidence of exchange if its *d*_*S*_ value was below the 2.5% quantile of the *d*_*S*_ of all nuclear genes, to have potentially transferred if *d*_*S*_ was between the 2.5% and 50% quantiles, and to show no evidence of transferring if above the 50% quantile.

We also examined polymorphism within TE families for evidence of horizontal transfer. We estimated Tajima’s *D* of each TE using *Popoolation* (Kofler *et al*. 2011),with the TE copy number as the sample size. As negative Tajima’s D may reflect recent expansion of a TE family (Bartolomé *et al*. 2009). We compared the levels of polymorphism shared among TEs in each species between potentially transferred TEs (*d*_*S*_ < 2.5% quantile) and TEs that are unlikely to have transferred (*d*_*S*_ > 50% quantile).

### Detecting long range horizontal transfer events with other Drosophila species

We attempt to identify long range transfers from other *Drosophila* species. To do this, we separated all known *Drosophila* TEs by their super families, including our set of *D. pseudoobscura* group TEs, and aligned the TE sequences within each superfamily using *MAFFT* (Katoh *et al*. 2002) and generated phylogenies for these using *PhyML* (Guindon *et al*. 2010). We then extracted patristic distance matrices for each superfamily using *Patristic* (Fourmant and Gibbs 2006) and compared each distance to the nuclear genome comparison performed previously for these genomes (Chen *et al*. 2014).

## Results and Discussion

### Transposable element annotation of the D. pseudoobscura group genomes

We identified insertions of the 157 well-supported TE families in the reference genome of the five species, and assessed their TE content using four metrics: the proportion of the reference genome masked (using *RepeatMasker* (Tarailo-Graovac and Chen 2009)), the proportion of short reads mapping to each TE sequences, the number of insertions in each genome using short read data (using *PopoolationTE2* (Kofler *et al*. 2016b), demonstrated across genomes in Supplementary Figure 2) and the estimated copy number of each TE family (Table 1 and Table S2). We also estimated the density of TE content across the genome (in masked bp/Mbp) using the proportion of the reference masked by *Repeatmasker*.

**Table 1:** Number of TE families and counts by order in species. % of reads mapping to each order in each species, number of copies found based on coverage relative to chromosome 3, % of the reference genome masked by each order for each species and number of insertions found using PopoolationTE2 (Kofler *et al*. 2016a). As LTR elements often exist not as complete insertions, but as solo-LTRs resulting from illegitimate recombination, coverage for the LTR elements was estimated for both solo LTRs and LTR bodies separately, with the average taken across the combined sequences. We tested for extrachromosomal circular DNAs such as from Helitrons and Polintons via comparisons between copy numbers and insertion numbers. We excluded the unknown families from the total insertion counts.

|  |  | Reads |  |  | Reference | PPTE2 |
| --- | --- | --- | --- | --- | --- | --- |
| Species | TE Order | No. families | % reads | est. num | % genome | est. num |
| ***D. pseudoobscura*** | TIR | 31 | 1.745 | 414 | 0.98 | 292 |
|  | LTR | 72 | 8.875 | 2230 | 7.21 | 1846 |
|  | LINE | 35 | 3.633 | 1121 | 2.85 | 927 |
|  | RC | 3 | 1.852 | 978 | 1.21 | 978 |
|  | Polinton | 1 | 0.417 | 149 | 0.081 | 29 |
|  | Unknown | 2 | 0.332 | 22 | 0.017 | 6 |
|  | Total * | 142 | 16.522 | 4892 | 12.33 | 4072 |
|  | Total (including Unknown) | 144 | 16.854 | 4914 | 12.5 | 4078 |
| ***D. persimilis*** | TIR | 31 | 1.547 | 413 | 1.29 | 392 |
|  | LTR | 72 | 14.273 | 2260 | 12.95 | 1919 |
|  | LINE | 35 | 6.956 | 1301 | 5.76 | 958 |
|  | RC | 3 | 4.43 | 1781 | 3.41 | 1755 |
|  | Polinton | 1 | 0.034 | 46 | 0.18 | 46 |
|  | Unknown | 2 | 0.543 | 76 | 0.025 | 7 |
|  | Total * | 142 | 27.24 | 5801 | 23.59 | 5070 |
|  | Total (including Unknown) | 144 | 27.78 | 5877 | 23.615 | 5077 |
| ***D. miranda*** | TIR | 31 | 0.892 | 262 | 0.87 | 258 |
|  | LTR | 67 | 7.19 | 973 | 2.21 | 925 |
|  | LINE | 36 | 5.367 | 1431 | 1.25 | 1059 |
|  | RC | 5 | 1.484 | 1934 | 1.16 | 1934 |
|  | Polinton | 1 | 0.054 | 9 | 0.024 | 9 |
|  | Unknown | 2 | 0.337 | 4 | 0.015 | 4 |
|  | Total * | 140 | 14.987 | 4609 | 5.51 | 4185 |
|  | Total<br>(including<br>Unknown) | 142 | 15.324 | 4613 | 5.525 | 4189 |
| ***D. loweii*** | TIR | 31 | 1.396 | 495 | 0.382 | 381 |
|  | LTR | 74 | 6.883 | 1366 | 1.55 | 740 |
|  | LINE | 34 | 3.839 | 933 | 0.799 | 449 |
|  | RC | 5 | 1.245 | 813 | 0.363 | 523 |
|  | Polinton | 1 | 0.054 | 7 | 0.013 | 7 |
|  | Unknown | 9 | 0.641 | 265 | 0.087 | 241 |
|  | Total * | 145 | 13.417 | 3614 | 3.1 | 2100 |
|  | Total<br>(including<br>Unknown) | 154 | 14.058 | 3879 | 3.187 | 2341 |
| ***D. affinis*** | TIR | 9 | 0.872 | 278 | 0.177 | 230 |
|  | LTR | 47 | 4.328 | 630 | 1.427 | 832 |
|  | LINE | 13 | 5.223 | 530 | 0.406 | 339 |
|  | RC | 4 | 1.351 | 369 | 0.245 | 369 |
|  | Polinton | 1 | 0.068 | 35 | 0.041 | 35 |
|  | Unknown | 10 | 1.192 | 206 | 0.098 | 206 |
|  | Total * | 74 | 11.842 | 1842 | 2.29 | 1805 |
|  | Total<br>(including<br>Unknown) | 84 | 13.034 | 2048 | 2.394 | 2011 |

**Figure 2:**
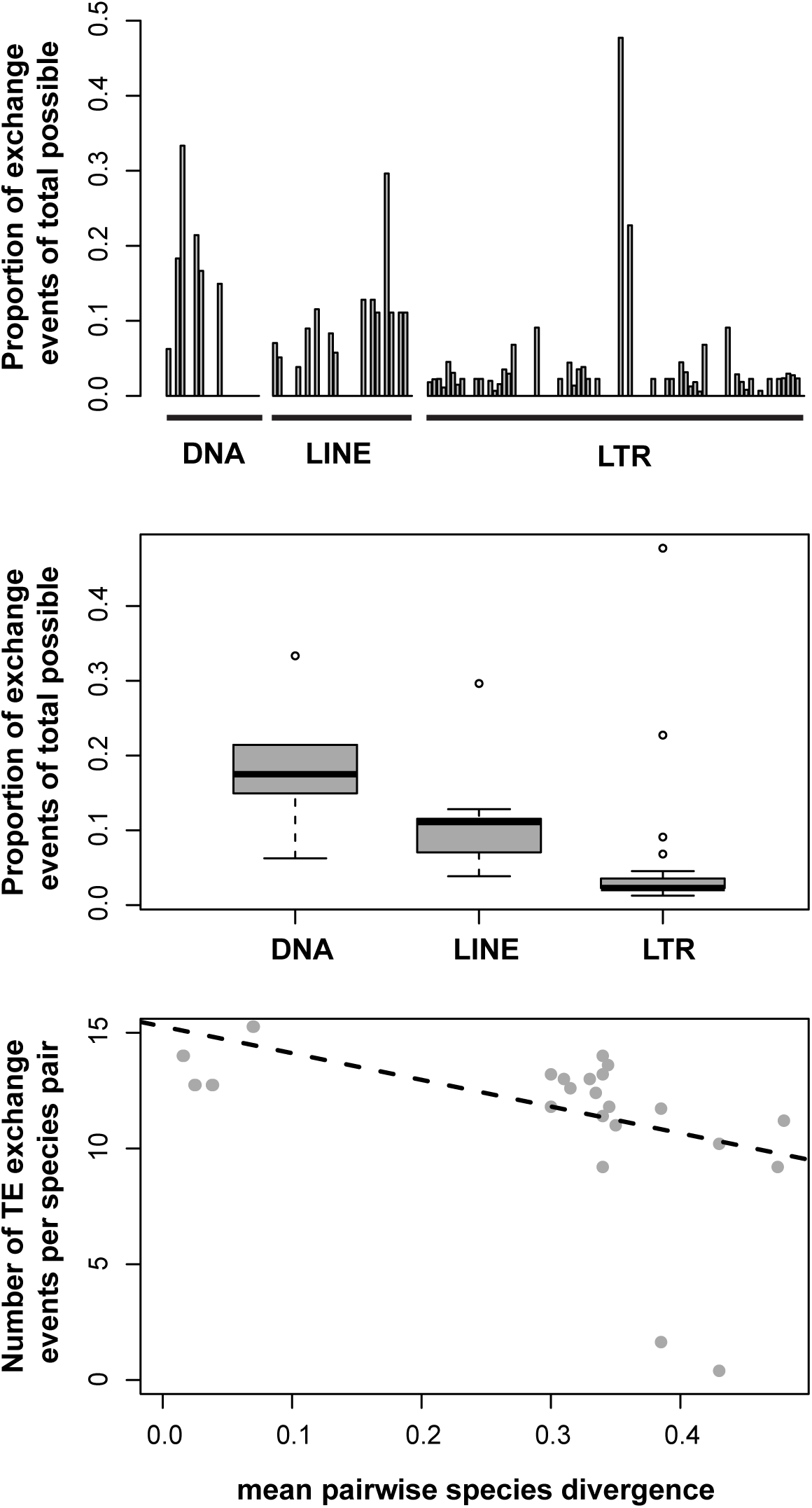
**A**. Each TE family and the proportion of times they show a lower divergence value than the mean divergence between the host species. **B**. Of the transferring TEs, the proportion of times these TEs are exchanging, grouped by TE order. **C**. Comparisons between the proportion of exchange events and the pairwise divergence between species, for exchanged TEs.

Across all species, for all measures of TE content we find a significant linear correlation between measures (Table S2, Spearman’s Rank Correlation *p*-value < 0.00213), though the strength of the correlation is weak for all species between the proportion of the genome masked at the family level versus the copy number of the TE family, and the insertion count versus the proportion of the genome masked (ρ < 0.58), suggesting that the proportion of the genome masked may be an inconsistent measure of TE density. As expected, correlations between measures of TE content in the species with genomes assembled only from short reads are lower (Table 1, Table S2) (Hoen *et al*. 2015; Rius *et al*. 2016), and the estimates of TE content for these species are likely more unreliable. We therefore limit our analysis of TE content to the two genomes with available long read data, *D. pseudoobscura* and *D. persimilis*. We also identified 15 sequences that pass all filters, but cannot be assigned to a TE order, we have included these sequences and their statistics, but have not included these sequences in further analyses (e.g. the 2 unknown sequences in *D. pseudoobscura*, Table 1, Table S2).

Because the *D. pseudoobscura* and *D. persimilis* genomes were originally assembled from long reads (and the *D. pseudoobscura* genome has also been assembled with the help of PacBio information and Sanger sequence information) (Richards *et al*. 2005; Clark *et al*. 2007), the TE content of these two species is already well-annotated. We found 116 previously identified *pseudoobscura* TE families using our pipeline. We also found two TE families from other species, and 28 additional putative TE families that passed all our filters in these two species. For *D. pseudoobscura*, we were able to use RNAseq data from (Duff *et al*. 2015) to determine whether these elements showed evidence of expression in embryos, using publically available expression data. We estimated RPKM for both novel and known TEs from these data; of the novel TEs, nine had appreciable levels of expression (Figure S3, FPKM > 1), a similar proportion to that of the known TEs (49 of 116). Similarly, we used sequences from embryonic small RNAs to ask if suppressive small RNAs are produced against these TE families. We extracted TEs with at least 20 small RNAs mapping to them, which comprised 114 of the 116 known TEs and all 28 of the novel TE sequences (Table S7 & 8). Most of these elements (108 of 140) had piRNAs generated against them (using the 24-29nt range generally used to identify piRNAs in other species (Ghildyal and Zamore 2009), and 27 elements also had homologous siRNAs (21-23nt small RNAs) (Figure S3, Table S2) (Obbard *et al*. 2009). A subset of the piRNAs, those produced in the germline (Aravin *et al*. 2007), are expected to show signatures of “ping-pong” amplification— small RNAs that match both sense and anti-sense strands of the TE sequence, an enrichment of these that show a 10bp overlap, a uracil in position 1 of sense strand piRNAs, and adenosine in position 10 of the antisense small RNAs) (Aravin *et al*. 2007). We found that 60 elements (53 known families and 7 novel; 36 LTRs, 15 LINEs, 7 DNA transposons & 2 helitrons) showed signatures of ping-pong amplification— from inspecting *Sequence Logos* plots (Table S2) (Schneider and Stephens 1990); novel and known elements showed ping-pong small RNAs at similar rates (Figure S3, Mann-Whitney U test W = 24, p > 0.1676). As these TE sequences are all multicopy, these measures of expression are mainly useful to show that the putative novel TEs have characteristics like those of the known TE sequences (Mann-Whitney U test W = 37, p > 0.05, Figure S3).

In all, we found 12.33% and 23.59% make up the reference genomes of these species, (Table 1). In contrast to a previous study, which found similar proportions of LTRs and LINEs in the *D. pseudoobscura* genome (Clark *et al*. 2007), we find over twice as much TE content due to LTR vs. LINE retrotransposons (Table 1); it is worth noting an additional effort was put into finding novel LTRs in the putative TE set using *LTRHarvest* (Ellinghaus *et al*. 2008).

In the remaining species, we find 20 additional families not found in *D. pseudoobscura* and *D. persimilis* (Figure 1B). The 57 TE families shared among all five species constitute most of the TE content (73-84% of insertions and 53-78% of each species reference TE content, Table S2), but vary in copy number between species (e.g. HelitronN-1 in *D. miranda* and *D. lowei*, Table S2), possibly due to stochastic expansion and loss of families over time. For example, we find *HelitronN-1_DPe* has 1927 insertions and makes up 1.1% of the genome of *D. miranda*, while it has only 727 insertions in *D. lowei*, comprising of 0.14% of the genome (Table 1, Table S2). This is likely due to the collapsing of *Helitron-1* and the closely related ISX sequence that has been co-opted for dosage compensation in *D. miranda* (Ellison and Bachtrog 2013). These differences can be further seen in the distributions of copy numbers in families, which differ between species (Figure S3, S4).

While we annotate the *D. miranda, D. lowei* and *D. affinis* genomes using a pipeline identical to that for other species, we suspect we have underestimated the TE content of these species. There are three main reasons. First, the genome assemblies for these species rely on short reads (Palmieri *et al*. 2014), which can lead to under-representation of the TE content of the genome (Rius *et al*. 2016). Similarly, previous estimates of the TE content of *D. pseudoobscura* and *D. persimilis* were much lower likely due to more fragmented genomes (Clark *et al*. 2007). Second, we may have missed TE families unique to these species, or may have recovered them only as fragments, as it is easier to recover full-length TE sequences if they closely match sequences already in RepBase as is true for the *D. persimilis* and *D. pseudoobscura* sequences. Finally, and likely the most important reason, the material sequenced for the reference genomes for *D. lowei, D. miranda* and *D. affinis* was adult females and not mixed sex embryos as for the others (Richards *et al*. 2005; Clark *et al*. 2007). Thus the other genomes contain the TE-rich Y-chromosome, which appears to be cytologically quite large in these species (Dobzhansky 1935, 1937), and may shows less under-replication of the TE-rich heterochromatin than adult samples.

In *D. persimilis* we found the same TE families as in *D. pseudoobscura*, but estimated 23.59% of the *D. persimilis* reference to be repetitive content versus 12.33% in *D. pseudoobscura*, implying 21.3MB more repetitive content in the *D. persimilis* reference genome compared to *D. pseudoobscura*. Previous annotation from the 12-genomes project found lower TE content as a proportion of the genome than that found here (3% and 8% *vs*. 12.33% and 23.59% here), but a similar ∼2-fold enrichment in TEs for *D. persimilis* (Clark *et al*. 2007). While it is true that *D. persimilis* has a larger genome than *D. pseudoobscura* ((Gregory 2005), the two species genomes are estimated to differ only by ∼2Mb (Bosco *et al*. 2007; Gregory and Johnston 2008).

The higher TE content of *D. persimilis* is not due to the presence of additional families, as the same families occur in both species (Table 1, Figure 1). In fact, as these species hybridize occasionally (Noor *et al*. 2007), it would be surprising if their TE families remained very distinct. Estimates of copy number from coverage of short read data (collected from adult females in both species) shows more copies of each TE family in *D. persimilis* than *D. pseudoobscura* (46.8 vs. 39.6 on average), but the difference is highly non-significant (Mann-Whitney U, p = 0.669).

Simple coverage differences of TEs, could, in principle, be explained by differences in under-replication of TE between the strains or species. But this is unlikely, as the coverage differences are also consistent with genome-size estimates, insertion number recovered and the proportion of the genome repeat masked (significant associations between all, as stated previously). If the difference is genuine, it could be due to the differences seen in a few families with large numbers of insertions in *D. persimilis*, such as *Gypsy10_Dpse, HelitronN-1_Dpe, Gypsy17_Dpse*, and *MiniME_DP* (Table S2). Based on coverages of each TE sequence in each species, we estimate that *D. persimilis* has, at most, ∼5Mb more TE content than *D. pseudoobscura*, consistent with the minor differences in genome size found between the two species (Bosco *et al*. 2007), suggesting a large amount of genomic content is missing from the *D. pseudoobscura* reference genome.

It is possible that an accumulation of TEs in the fixed inversions between *D*.*pseudoobscura* and *D. persimilis* could explain the large difference in TE content, due to the reduced genomic exchange in these regions (Machado *et al*. 2007), allowing insertions to accumulate in one species but not the other. Consistent with this idea, we find that LTR retrotransposons are at significantly higher densities of TE insertions within these inverted regions in *D. persimilis* when compared to *D. pseudoobscura* and the uninverted regions (File S2, Insertions per MB, using inversion windows defined in (Avila *et al*. 2014); Mann Whitney U test: LTR inside inversions W = 53686, p = 5.674e-05, LTR near inversions W = 16604, p-value = 0.1128 LTR outside inversions W = 290520, p-value = 0.1407). However, we find RC and LINE insertions are at significantly higher densities in *D. persimilis* regardless of genomic location (Insertions per MB, Mann Whitney U test: W > 335780, p-value < 0.0001303 for inside, outside and near inverted regions) and no difference in TIRs (W < 790, p-value > 0.37), suggesting that the fixed inversions are not the only explanation.

One final possibility, the Y chromosome of these species may also play a role in both the genome size and TE content differences of the species. While considerable variation exists in the size of *D. pseudoobscura* Y chromosome size across types (Dobzhansky 1935, 1937), the *D. persimilis* Y chromosomes are limited to the largest of these types (Types I, II and III). As the *D. pseudoobscura* reference genome was likely generated from a strain containing the smallest Y chromosome type (Standard/Arrowhead, likely type V) (Dobzhansky 1937; Dobzhansky and Sturtevant 1937), while the *D. persimilis* genome strain used to generate their genome likely contains the most common *D. persimilis* Y, the largest of the chromosome types (Dobzhansky 1937). Previous work has also found Y-linked variation in *D. melanogaster* and *D. simulans* to be associated with phenotypic variation in a number of factors including TE regulation, it is possible that a larger Y can cause poorer TE regulation, due to the increased heterochromatin load in the genome (Sackton and Hartl 2013; Francisco and Lemos 2014). This is possibly the case between *D. pseudoobscura* and *D. persimilis*, where the larger Y chromosome may have led to the poorer regulation of TE families, leading to the ∼5Mb expansion of TEs in *D. persimilis*.

### Several transposable element families show evidence of ancient horizontal spread between species

As a majority of TEs are likely acquired in a species by horizontal transfer by closely related species (Burt and Trivers 2006; Peccoud *et al*. 2017). We examined our set of TEs for evidence that they had been horizontally acquired from another *Drosophila* species by comparing the patristic distance of all *Repbase* TEs pairwise to the average patristic distance of *pseudoobscura* group TEs (Kohany *et al*. 2006), after building a phylogeny of each superfamily (Supplementary Figure 3).

**Figure 3:**
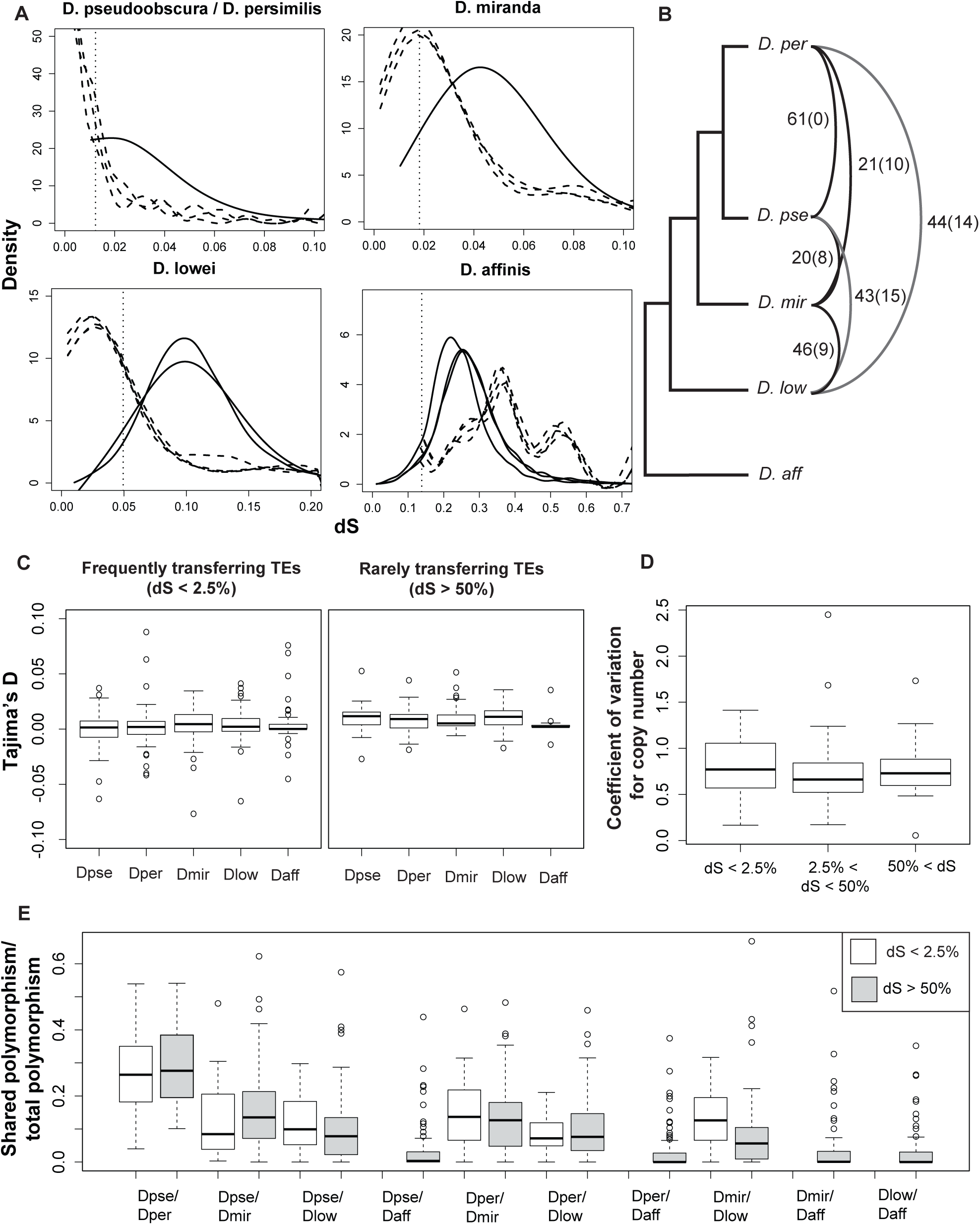
**A**. Pairwise comparison of silent site diversity (d_S_) for nuclear genes (solid line) and shared TEs (dashed lines) between *D. pseudoobscura, D. pseudoobscura bogotana, D. persimilis* and other species. The lower 2.5% quartile for nuclear d_S_ is shown as the dotted vertical line **B**. The number of transfer events for transposable elements based on d_S_, the number in brackets shows events that can be seen in the assembled phylogenies. Note that many events could be occurring between species vertically as well as horizontally. **C**. Comparison of Tajimas D across species for frequently exchanged TEs and rarely exchanged TEs shows no difference, suggesting no population expansion. **D**. No more variation in copy number of rarely exchanged TEs than with frequently exchanged TEs. **E**. Proportion of shared nucleotide polymorphism sites between TE sequences in species, out of total nucleotide polymorphism sites, divided by TE families with low Ks relative to nuclear genes and TEs with higher d_S_

Across 4096 pairwise comparisons, we found 230 where the TE patristic distance was lower than the previously found genic distance (Table S7) (Chen *et al*. 2014). These events were limited to 63 of 157 TEs, with most these TEs having lower patristic distances than entire species groups (such as the *D. rhopaloa/D*.*elegans/D*.*ficusphila* species subgroup), consistent with a transfer of the TE between the common ancestor of the species and the *D. pseudoobscura* group, followed by a diversification into multiple elements seen today. 42 of these transferred elements are LTRs, 6 are DNA transposons and 15 are LINEs (Table S7, Figure 2A). While a higher proportion of LTRs are transferred between species, each of these families only appears to have been exchanged with a single species, rather than multiple, likely because of the recent expansion of LTRs in *Drosophila*, compared to the more ancient expansion of most LINEs and DNA transposons (Figure 2B). Among these transferred elements, we find a *piggyBac* element that was acquired from *D. busckii*, several elements from the Asian subgroup of the *D. melanogaster* group, (such as *P_226* with *D. elegans, Jockey_185* with *D. rhopaloa* and *I_149* with *D. ficusphila*) and several Jockey elements are closely related to elements found in the *Drosophila* clade species, such as with a *D. virilise* ancestor. We compared the proportion of TEs showing HT events between species to the patristic distance to each species, we find a significant negative linear correlation between the species genic patristic distance and the proportion of TEs (Figure 2C, Binomial GLM logistic regression, z-value = −7.88, p-value< 2e-16), agreeing with previous findings that horizontal acquisition is more likely between closely related species (Peccoud *et al*. 2017).

### Evidence of recent recurrent horizontal transfer between species

In the *D. melanogaster* group, in addition to occasional bouts of catastrophic invasion, many elements appear to have been transferred commonly between close relatives in the group (Daniels *et al*. 1990; Clark and Kidwell 1997; Bartolomé *et al*. 2009). For *pseudoobscura* group TEs found in at least 3 species which had previously described coding regions (101 TEs, 39 for comparisons to *D. affinis*), we compared the silent site divergence (*d*_*S*_) of TEs found between species to the *d*_*S*_ of host genes. Overall, we found a significant reduction in synonymous divergence relative to host genes for all comparisons (Mann-Witney U test *p-value* < 0.05), excluding those involving *D. affinis* (Figure 3A). We find 76 TE families below the 97.5% quantile of nuclear gene d_S_ in at least one comparison suggesting potentially recent transmission between species (51 of 62 LTRs, 19 of 30 LINEs and 6 of 8 DNA transposons). Inconsistent with horizontal transfer, there is not a depletion of non-LTR retrotransposon (LINE) elements found here.

We also compared the phylogenies of the TEs to that of the species, again looking for evidence of horizontal transfer. Of these families, 41 have phylogenies that differ from the species tree and group the two species with little divergence together. It is possible these differences are due to incomplete lineage sorting, or gene tree discordance, it is also possible that horizontal transfer has occurred for this family between these species, and so may support HT for 34 LTRs, 3 DNA transposons and 5 LINEs (All of which are below the 97.5% quantile for genomic *d*_*S*_, Figure 3B; Table S2). Again, we find no evidence of exchange with *D. affinis*. In the *D. pseudoobscura* subgroup each species can hybridize with others to some degree (though likely not occurring in nature; Machado et al. 2007), therefore, we cannot determine if these apparent transfer events are true horizontal events or vertical transfer via hybridisation. We do see slightly elevated proportions of TIRs & LTRs when comparing phylogenies, consistent with horizontal transfer as suggested previously in (Sánchez-Gracia *et al*. 2005; Bartolomé *et al*. 2009). Conversely, we found *d*_*S*_ between species and *D. affinis* was significantly higher for TEs than host genes, consistent with the allopatric separation limiting HT events seen between species and possibly unconstrained evolution in the TEs (Figure 3A, Mann Whitney U test: *p <* 3.5e-08, Table S5).

By comparing Tajima’s D for each TE in species we can look for strongly negative D, consistent with a copy number expansion following horizontal transmission (Bartolomé *et al*. 2009). All comparisons show equal levels of D in each species, close to 0, implying that each species already share the TE families, resulting in no expansion in copy number (Figure 3C). Consistent with this, we find all the TEs potentially shared between species have shared polymorphism (Figure 3E), which is not expected if acquisition is recent and purely horizontal. However, this result conflicts with our expectation from the nuclear d_S_ comparison, which we expect to be at similar levels to TE d_S_ if there is hybridization. This result suggests pervasive transmission between species, resulting in polymorphism being exchanged between species several times, rather than once, resulting in no excess of low frequency polymorphism (Tajima 1989; Bartolomé *et al*. 2009). Alternatively, there is less constraint on polymorphism in transposable elements, allowing polymorphisms to drift to higher frequencies in shorter periods of time following their horizontal acquisition.

Interestingly, 10 TE families appear to transfer between species in all comparisons (d_S_ < 0.25% quantile: 1 TIR, 1 LINE and 8 LTRs), while 21 show no evidence of transfer (d_S_ > 50% quantile: 1 TIR, 1 helitron, 11 LINEs and 9 LTRs), suggesting that rates of transmission are highly dependent on the TE family and its activity. We also see large differences in copy numbers of each family in each species. We next looked to see if a lack of exchange can lead to changes in copy number of a family and explain the differences between *D. persimilis* and others.

We compared changes in copy number over all the species (via the coefficient of variation), for pervasively transferring TEs, non-transferring TEs and all other TEs. We find no difference in the coefficient of variation of copy number for pervasively transferring families and non-transferring families (Figure 3D; Mann Whitney U *p*-value > 0.19 for all comparisons), suggesting that reduced transmission between species isn’t altering dynamics of the families compared to pervasively transferring families. This low divergence and no evidence of family expansion has two possible explanations: 1. There may be gene flow to some degree between these species in the wild, while the genes are likely not introgressed due to incompatibilities or lower fitness, their linked TEs will be transpose more readily after hybridisation, becoming unlinked from this gene. This variant will then be maintained in the new host, resulting in reduced divergence for the TE family between the species. 2. Due to the sympatry of the *pseudoobscura* subgroup, there may have been recurrent horizontal transmission between species, for TE families already present in each of the species, resulting in the low *d*_*S*_, but shared polymorphism and lack of copy number expansion. The lower numbers of LINE families found exchanging between species supports the idea of horizontal exchange, however the supposed numbers of exchanges (up to 61) between species is unprecedented, giving more support to vertical exchange of TEs. Despite this pervasive TE exchange of some families, TE dynamics may be changing within the species, leading to the differences seen in TE families, densities and family copy numbers.

Like *D. melanogaster*, the *D. pseudoobscura* group shows highly active TEs that appear to be constantly undergoing a cycle of acquisition, expansion and high activity, suppression and finally extinction. Strangely, despite TE exchange between species, the group shows distinct differences in TE content and TE densities. Though some of these differences are due to differences in quality of assembly of each species genome and method used to identify TE insertions, we find a distinct expansion in TE numbers in *D. persimilis*. We find these differences are likely due to stochastic differences in expansion and extinction between shared families, and not due to differing activities in novel and private families compared to these shared families. Overall this suggests that despite frequent gene flow, TE dynamics can evolve rapidly due to stochastic factors across the lifetime of a family.

Due to the history of the first recorded instance of a horizontal transfer of a transposable element, we tend to think that these transfers are rare, likely catastrophic events. However, an expanding body of evidence suggests that these events are likely a common occurrence throughout genomes, becoming more and more common the more closely related two species are. This transfer of elements is possibly even recurrent in some cases and, due to the presence of closely related sequences within piRNA clusters, reducing the fitness costs of a transfer event, such as the invasion of P-element into *D. melanogaster*. Our results support the idea that TEs are highly fluid, moving between genomes easily without leading to the expansion of TE content in a species genome, or heavy catastrophic events such as was seen in laboratories with the invasion of P-element.

## Acknowledgements

Thanks to L. Endler and D. Gómez Sánchez for discussion and advice concerning the bioinformatic analysis. We are grateful for helpful discussion provided by K. Senti, R. Kofler, B. Charlesworth, A. Clark, R. Unckless and J. Blumenstiel. Thanks for S. W. Schaeffer and M. Noor for providing the data used in this survey and advice concerning how the data should be used. The *D. pseudoobscura* line sequencing was funded by a National Institute of Health (NIH) grant R01-GM098478 to S. W. Schaeffer. Funding for TH was provided by the Max Kade foundation.

---

**Figure S1:** Pipeline for TE annotation.

**Figure S2:** TE density across the genomes of each species, found using *PopoolationTE2*, sorted by TE order.

**Figure S3:** Comparison between putatively novel and known TE sequences for (A) length, (B) expression, (C) small RNA silencing expression and (D-F) copy number.

**Figure S4:** Distribution of TE copy numbers per species.

**Table S1:** *D. pseudoobscura* lines used in this study

**Table S2:** TEs found in *D. obscura* group. Sorted by if they are previously discovered or novel, then by Order and super family. Transmission states if the TE family is found to transfer between species

**Table S3:** Diagonal shows the total number of families found in each species for comparison.

**Table S4:** GLMs for three recombination maps versus TE accumulation, divided by order and super family. Done for both TE count (quasipoisson GLM) and TE density (binomial GLM). Significant values (*p*< 0.05) are shown in bold.

**Table S5:** For instances where no dS for nuclear comparisons are available, we used the dS between D. pseudoobscura and the species of interest.

**Table S6:** Number of unique and shared polymorphic sites for each species comparison, for each TE family.

**Table S7:** *D. pseudoobscura* TEs and the patristic distance from other TEs in their superfamily group, compared to the patristic distance between the TEs fly species and *D*.*pseudoobscura*

