## Supplementary Materials for "Extensive horizontal exchange of transposable elements in the *Drosophila pseudoobscura* group"

**Figure S1:** Pipeline for TE annotation.

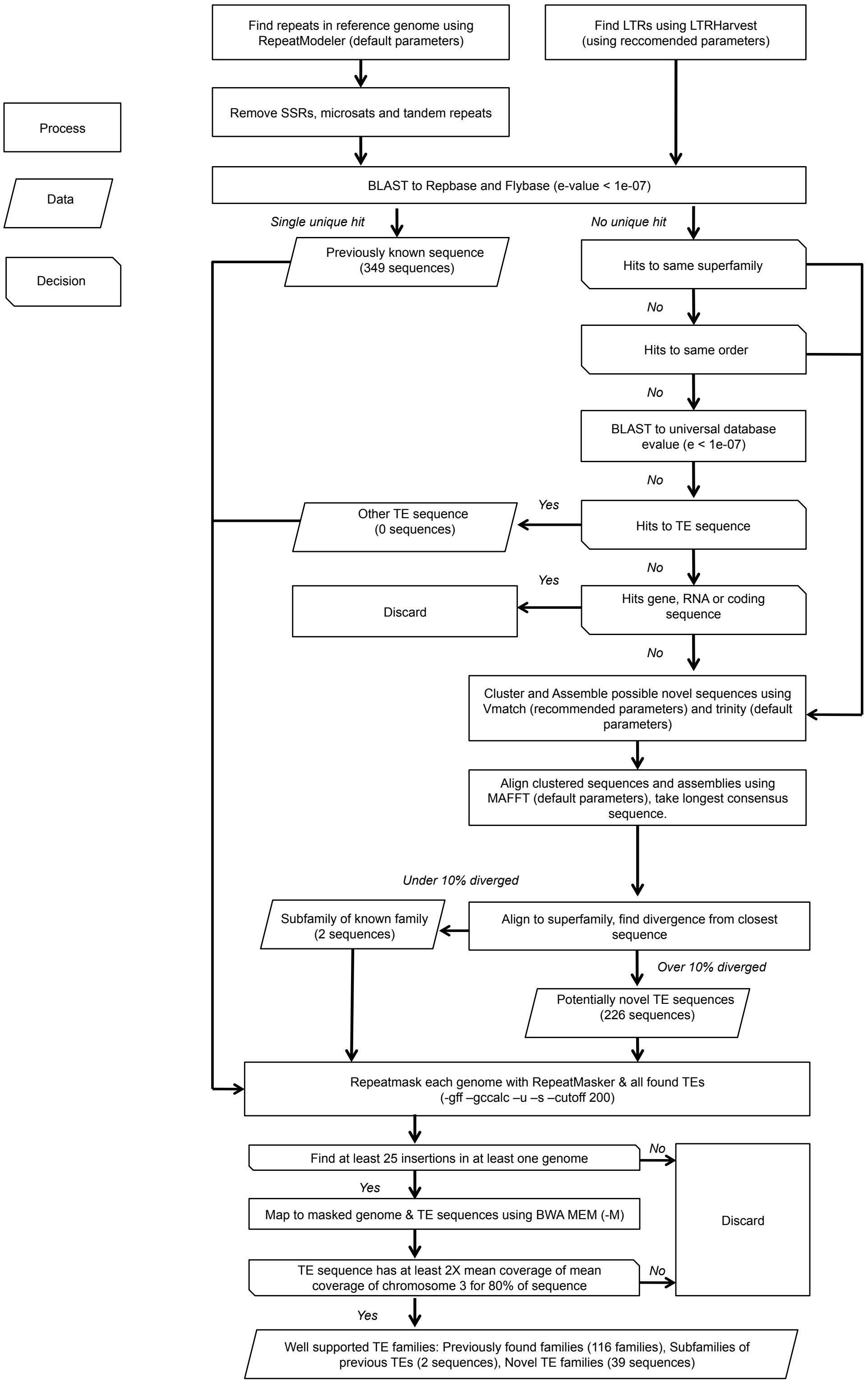

**Figure S2:** TE density across the genomes of each species, found using *PopoolationTE2,* sorted by TE order.

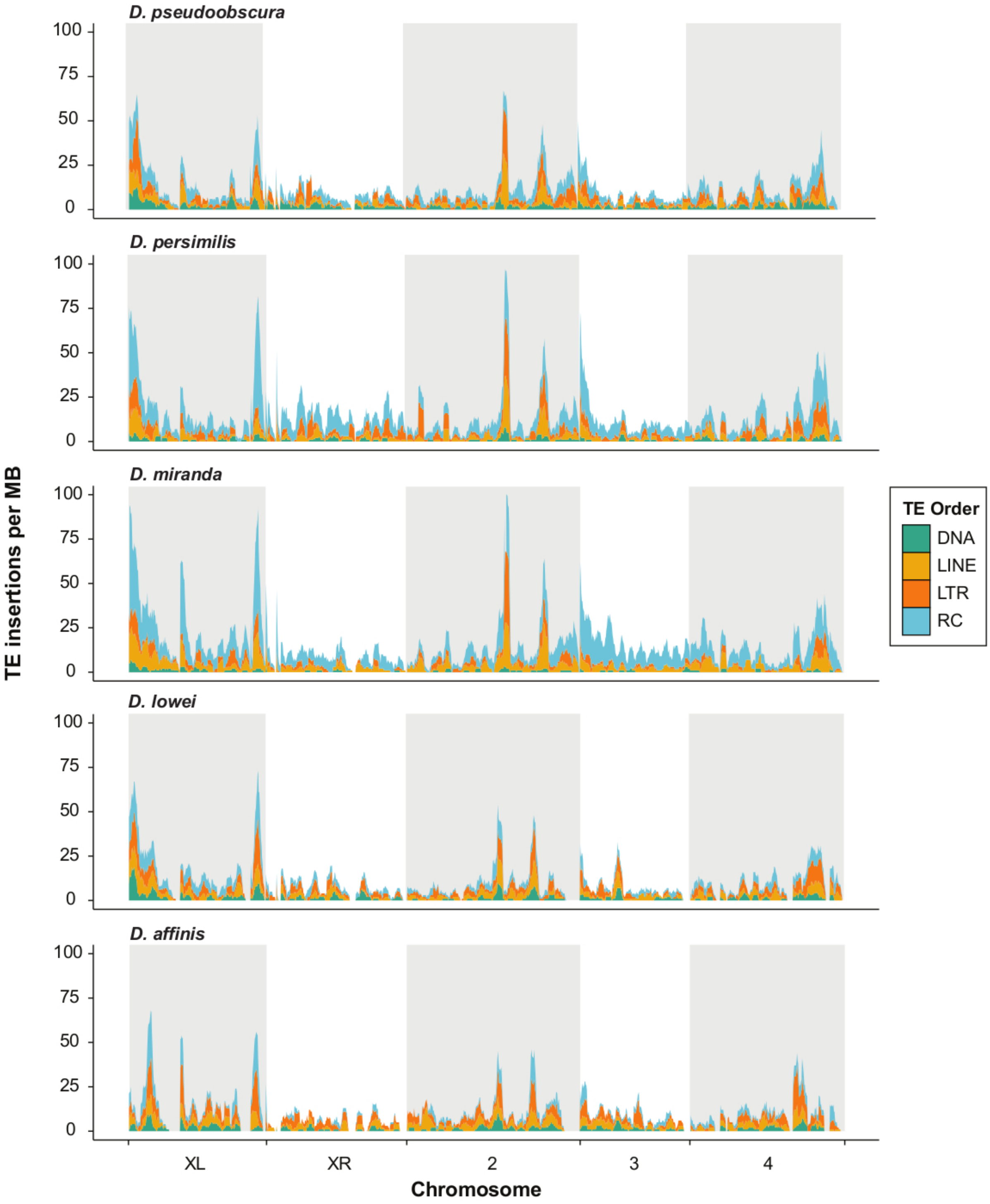

**Figure S3:** Comparison between putatively novel (grey) and known TE sequences (white) for (A) length, (B) expression, (C) small RNA silencing expression and (D-H) copy number.

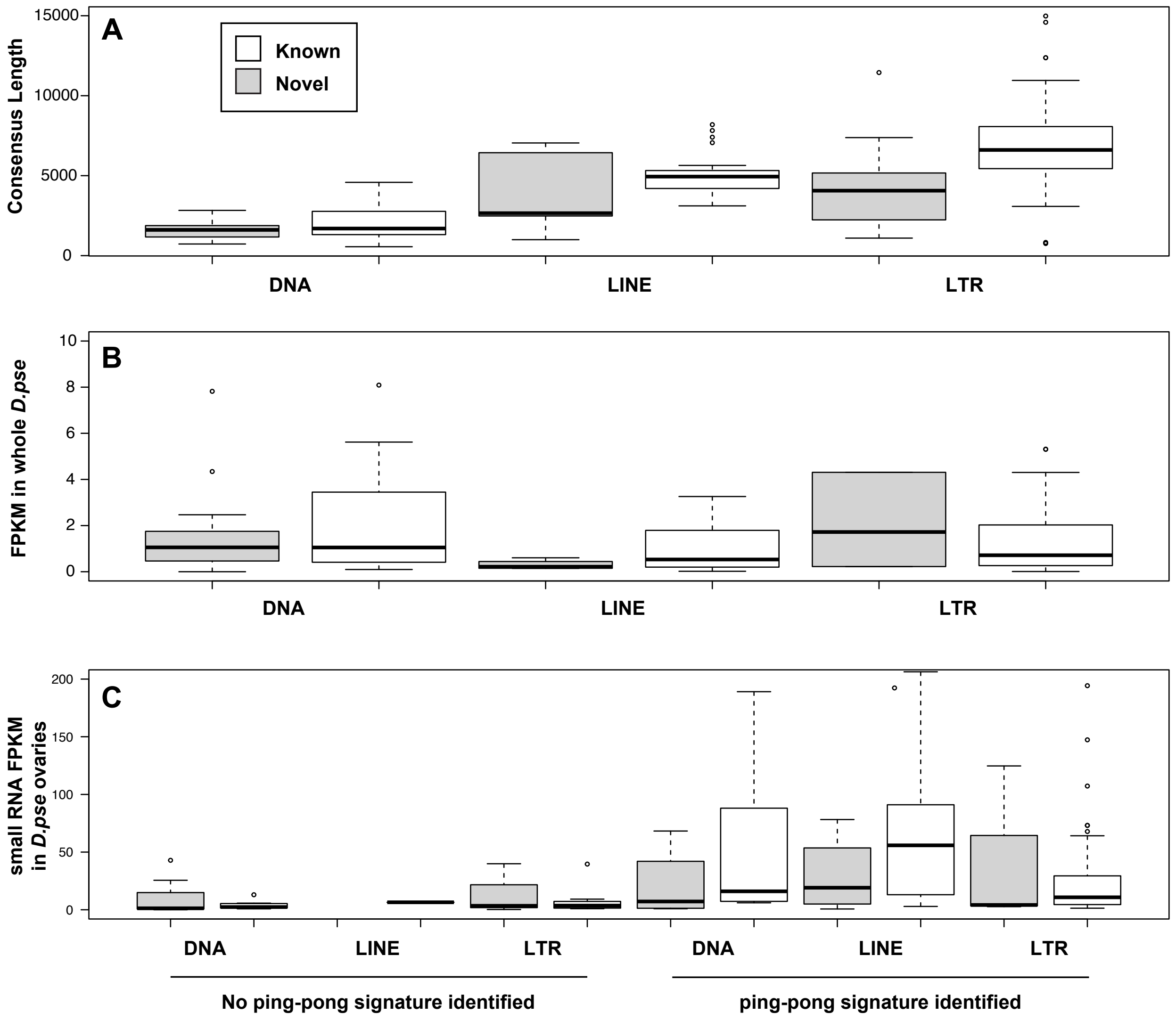

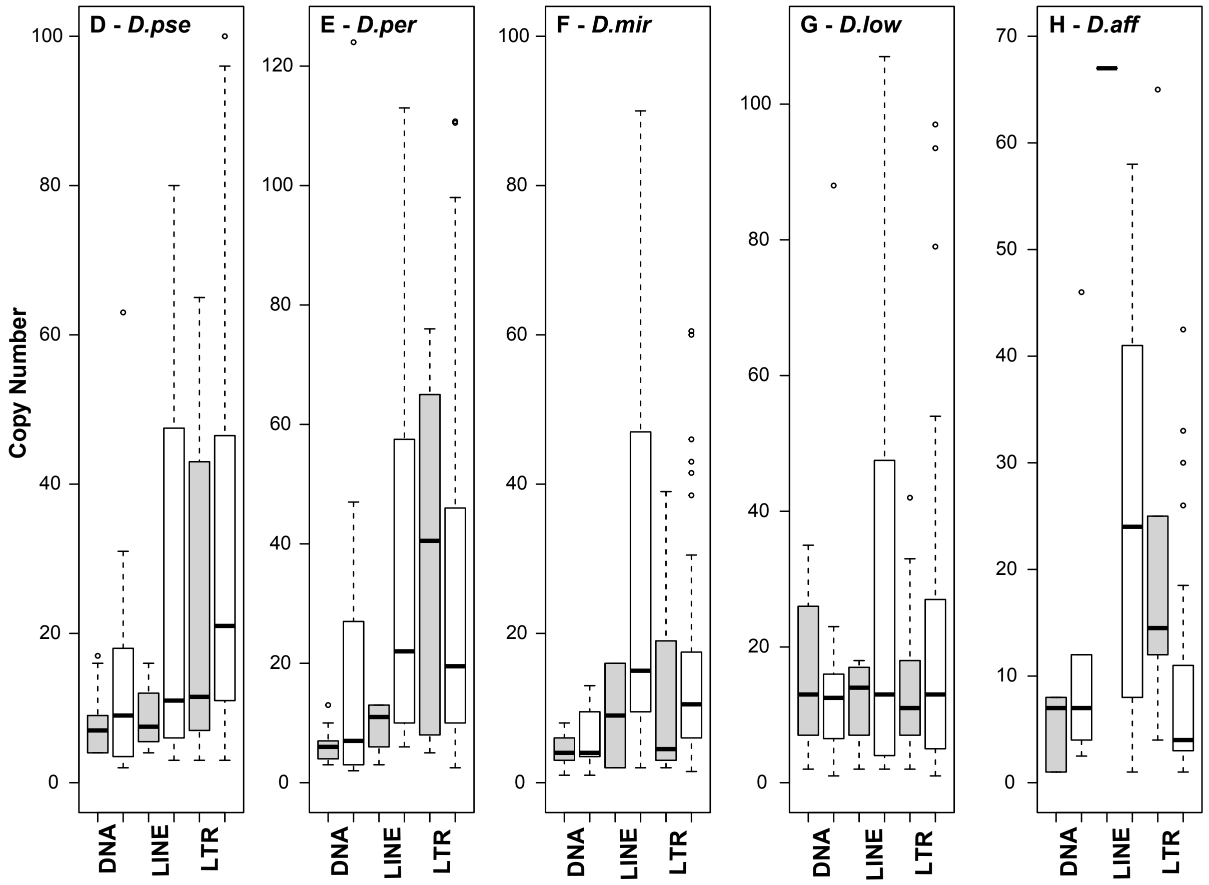
D - D.pse
E - D.per
F - D.mir
G - D.low
H - D.aff
Copy Number
DNA
LINE
LTR

**Figure S4: Distribution of TE copy numbers per species.**

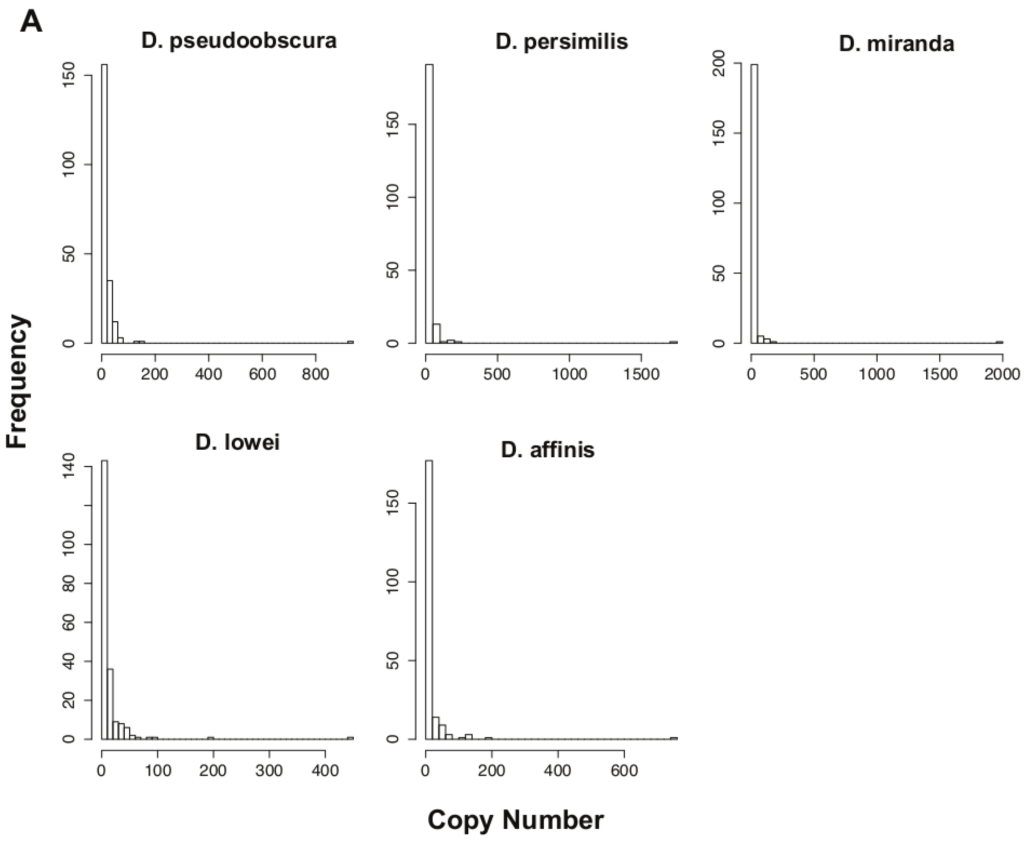
